# Chemogenetic activation of corticotropin-releasing factor-expressing neurons in the anterior bed nucleus of the stria terminalis reduces effortful motivation behaviors

**DOI:** 10.1101/2023.02.23.529717

**Authors:** Isabella Maita, Allyson Bazer, Kiyeon Chae, Amlaan Parida, Mikyle Mirza, Jillian Sucher, Mimi Phan, Tonia Liu, Pu Hu, Ria Soni, Troy A. Roepke, Benjamin A. Samuels

## Abstract

Corticotropin-releasing factor (CRF) in the anterior bed nucleus of the stria terminalis (aBNST) is associated with chronic stress and avoidance behavior. However, CRF+ BNST neurons project to reward- and motivation-related brain regions, suggesting a potential role in motivated behavior. We used chemogenetics to selectively activate CRF+ aBNST neurons in male and female CRF-ires-Cre mice during an effort-related choice task and a concurrent choice task. In both tasks, mice were given the option either to exert effort for high value rewards or to choose freely available low value rewards. Acute chemogenetic activation of CRF+ aBNST neurons reduced barrier climbing for a high value reward in the effort-related choice task in both males and females. Furthermore, acute activation of CRF+ aBNST neurons also reduced effortful lever pressing in high-performing males in the concurrent choice task. These data suggest a novel role for CRF+ aBNST neurons in effort-based decision and motivated behavior.

## Introduction

Chronic exposure to stressful experiences can precipitate mood disorders. In rodents, decades of work demonstrate that chronic stress exposure results in long-lasting changes to limbic circuitry and increased avoidance behaviors in tasks historically associated with anxiety or depression. However, deficits in reward and motivation behaviors are observed in several psychiatric disorders (including mood disorders) [1, 2]. Importantly, similar versions of these complex reward and motivation behavioral tasks can be performed in both rodents and humans, with analogous data analyses and interpretations [1]. These cross-species behavioral measures should provide enhanced translational validity and better guide therapeutic development than traditional avoidance behaviors. Importantly, humans diagnosed with mood disorders [2] and rodents exposed to chronic stress [3–5] exhibit reduced motivation to exert effort for rewards.

The bed nucleus of the stria terminalis (BNST) is a limbic region associated with chronic stress and avoidance behaviors. Exposure to unpredictable fearful stimuli activates the BNST in rodents [6] and humans [7–11]. BNST threat responsivity is increased in subjects diagnosed with post-traumatic stress disorder (PTSD) and generalized anxiety disorder [12–14]. The anterior BNST (aBNST) contains a subpopulation of GABAergic neurons that express corticotropin-releasing factor (CRF, also known as corticotropin-releasing hormone or CRH). Optogenetic activation of CRF+ aBNST neurons in rodents increases avoidance behavior [15–17], and stress-induced activation of CRF-expressing (CRF+) aBNST neurons is necessary and sufficient for the ensuing increases in avoidance (reviewed by [18]). Stress exposure produces long lasting changes in CRF+ aBNST neurons, such as increased PKA [16] and CRF signaling [16, 19, 20], as well as increased neuronal excitability [16, 19, 21–23]. Suppressing CRF signaling in the aBNST via PKA antagonism [16] or CRF antagonism [24, 25], or inhibiting CRF+ aBNST neurons with optogenetics [26] ameliorates stress-induced avoidance behaviors.

While CRF+ aBNST neurons promote avoidance, neural circuit connections and behavioral evidence suggest that CRF+ aBNST neurons may also be involved in stress-induced changes to motivation behaviors. CRF+ aBNST neurons project to [27–29] and inhibit [17, 30] the VTA and nucleus accumbens (NAc) shell. Stress increases CRF release onto the VTA [31], which reduces dopamine release [32] in response to rewards [33]. CRF+ aBNST neurons are also associated with the aversive effects of withdrawal [34], and promote drug-seeking, including reinstatement of cocaine seeking [35, 36] and cocaine-induced dopamine release [30]. In a two-choice progressive ratio task for sucrose alone or paired optogenetic stimulation of CRF+ BNST neurons with sucrose, rats lever pressed significantly less for the paired sucrose and laser stimulation [37]. These results add to evidence that activation of CRF+ aBNST neurons is aversive [17, 26, 38], but do not determine whether these neurons play a role in effortful behaviors.

Therefore, it remains unclear whether CRF+ aBNST neurons play a role in motivation behavior, such as willingness to exert effort to receive high value rewards. Since chronic stress in mice decreases effortful behaviors and results in long-lasting changes to BNST CRF+ neurons, including increased excitability in CRF-expressing neurons [16, 19, 21, 22, 39], we used chemogenetics to dissect the role of CRF+ BNST neurons in effortful motivation behaviors.

## Materials and Methods

### Transgenic and Wildtype Mice

All experiments were carried out in accordance with the National Institutes of Health guidelines for animal care and with approval from the Rutgers Institutional Animal Care and Use Committee. Mice were group housed and allowed access to ad libitum standard lab chow (Lab Diet 5001 Rodent Diet, Lab Diet, St. Louis, MO) and water, except when fasting during behavioral experiments. Throughout effort-related choice, concurrent choice, and free feed training and testing, animals were maintained at 85-90% of their ad libitum body weight via a daily supply of ~3 g food per mouse comprised of reinforcer pellets and/or supplementation with standard lab chow. Mice were maintained at a 12:12 light/dark schedule, with lights on at 6 AM, and all training and testing was completed between the hours of 7 AM and 4 PM.

A combined total of 26 (12 male and 14 female) C57BL/6J and 39 (20 male and 19 female) CRF-ires-Cre transgenic mice from Jackson Labs (Bar Harbor, ME, USA) were used. Heterozygous CRF-ires-Cre transgenic mice were produced by crossing C57BL/6J wildtype mice (WT, Jackson Laboratory #000664) with homozygous *Crf-ires-Cre* mice (Jackson Laboratory #012704), a strain generated by Taniguchi et al., 2011 [40] that expresses Cre recombinase in neurons with endogenous CRF expression. 18 (8 males and 10 females) heterozygous CRF-ires-Cre and 26 WT C57BL/6J (12 males and 14 females) underwent effort-related choice testing and 21 (12 males and 9 females) heterozygous CRF-ires-Cre mice underwent concurrent choice testing.

### Stereotaxic Surgery and Viral Injections

All transgenic mice (6-9 weeks) heterozygous for CRF-ires-Cre underwent a stereotaxic surgical procedure to induce virally mediated expression of Gq-coupled Designer Receptors Exclusively Activated by Designer Drugs (DREADDs) or a control virus. Mice were deeply anesthetized with 1-1.5% isoflurane and head-fixed in a stereotaxic apparatus (Kopf Instruments, Tujunga, CA, USA). Lidocaine (Lidoject, Covetrus) was topically injected, and skin was prepared with alternating application of betadine and 10% ethanol. Mice were bilaterally infused with 300 nl of virus into the anterodorsolateral BNST at the following coordinates: +0.14 mm anterior to Bregma; +/-1.00 lateral to the midline; −3.60 mm ventral from the surface of the brain, which resulted in viral expression throughout the aBNST. DREADD mice were injected with AAV9-hSyn-hM3D(Gq)-mCherry (AddGene Plasmid #, Watertown, MA, USA), virus titer 1.8 x 10^13^. Control mice were injected with one of two control viruses: AAV9-hSyn-DIO-mCherry (AddGene Plasmid #50459), virus titer 2.17 x 10^13^ or AAV9-hSyn.HI.eGFP-Cre-WPRE-SV40, virus titer 2.7 x 10^13^ (AddGene Plasmid #105540). The skull was sealed with bone wax (Med Vet International, Mettawa, IL, USA), the scalp was closed with VetBond (3M, St. Paul, MN, USA) and mice were allowed 3 weeks for recovery and viral expression before beginning training in either effort-related choice or concurrent choice.

### Clozapine-N-Oxide (CNO)

CNO (Sigma-Aldrich, St. Louis, MO, USA) was dissolved in 1 mL dimethyl sulfoxide (DMSO, Fisher Scientific) and adjusted to a final concentration of 0.5 mg/ml CNO in 7.5% v/v DMSO in sterile saline, then filtered through a syringe filter. Vehicle solution consisted of 7.5% v/v DMSO in sterile saline. 30 minutes prior to behavioral testing, the abdomen was prepared with a 10% ethanol wipe, then either 2 mg/kg CNO solution or Vehicle solution were injected intraperitoneally (i.p.).

### Effort-Related Choice

A cohort of 18 6- to 7-week-old CRF-ires-Cre mice (8 males and 10 females) underwent stereotaxic surgery and Y-maze barrier testing. A second cohort of 26 DREADD-free WT controls (12 males and 14 females) also underwent Y-maze barrier testing. The effort-related choice apparatus consisted of a Y-shaped maze composed of white Plexiglas. The Y-maze arms were 26 cm in length, with a uniform width of 7 cm and height of 20 cm. To clean and remove odors from the maze, the maze was sprayed with 70% ethanol between trials and allowed to dry. First, mice were trained to choose a high reward (HR) arm that contained four highly palatable 20 mg dust-free reinforcer pellets (Bio-Serv, Flemington, NJ, USA) over a low reward (LR) arm that contained only two reinforcer pellets. Then, effort-related choice was tested by placing a barrier in the HR arm. The HR arm was counterbalanced across animals, so 50% of subjects were assigned the HR in the left arm of the Y-maze. In all training and testing trials, mice were placed in a start box at the end of the Y-maze, then allowed 60 seconds to consume a reward in either the HR or LR arm. Mice were grouped so there was an average of a 5-minute inter-trial interval. Throughout training and testing, arm choice and latency to choose an arm were recorded by independent researchers blinded to treatment groups.

Training: After 3 weeks of recovery from surgery, mice were fasted to 85-90% of their ad libitum body weight. Mice first underwent two habituation trials where they explored the Y-maze for 15 minutes and consumed pellets in both the HR and LR arms. Next were three days of forced choice training, where mice were forced to select arms in 10 alternating trials by blocking either the HR or LR arm with a Plexiglas wall. Free-choice training sessions came next, where mice underwent 2 alternating forced choice trials and then 10 free-choice trials during which mice were allowed 60 seconds to consume the reward in either the HR or LR arm. All mice achieved a criterion of selecting the HR arm in 70% of trials after 16 days of free-choice training. From days 1-14 of free-choice training mice did not receive injections. However, on days 15 and 16, all mice were i.p. injected with saline 30 minutes prior to free-choice training to habituate them to the handling and injections. WT DREADD-free controls did not undergo stereotaxic surgery or viral injection, but they underwent the same Y-maze training protocol. They also reached 70% HR arm choice criteria after 16 free-choice training days, at which point all mice were injected with saline 30 minutes prior to one final session of free-choice training on day 17.

*Testing:* During all barrier testing sessions, a wire mesh barrier was placed into the HR arm. Mice then needed to physically climb the barrier to access the high reward. Barrier testing sessions consisted of 2-4 alternating forced-choice trials followed by 10 free-choice trials. Mice underwent 3 sessions of barrier testing at each barrier height: 10 cm, 15 cm, and 20 cm. Arm choice over the ten daily trials were recorded. After completion of 9 days of barrier testing, mice then underwent 3 discrimination sessions of 10 free-choice trials where 10 cm barriers were placed in both the HR and LR arms. All CRF-ires-Cre mice were injected 30 minutes prior to testing sessions with CNO. DREADD-free WT control mice were randomly assigned into groups that received either CNO or Vehicle injections throughout testing.

### Concurrent Choice Task

A cohort of 21 6–7-week-old CRF-ires-Cre mice (12 male and 9 female) underwent stereotaxic surgery and concurrent choice training and testing. Standard mouse operant chambers (Med Associates, Fairfax, VT) were housed in sound-attenuating cubicles inside a behavioral testing room. Operant chambers included two retractable response levers on one wall and a food delivery port connected to pellet hoppers dispensing reinforcer pellets. Training and testing consisted of 30-minute daily sessions, 5 days a week. During training, mice did not receive CNO or Vehicle injections.

Training: Concurrent choice began with 10 days of habituation to lever pressing using a fixed ratio (FR)1 reinforcement schedule. Mice then underwent training on FR1, variable ratio (VR)2, VR5, VR10, and VR20 schedules. Next came 10 days of progressive ratio (PR) baseline training sessions, where mice were required to press at increasing rates to receive a reward on a FR(n) x 15 schedule, wherein n increased by one for every 15 reinforcers received. Mice then began PR with concurrent chow (PR/CHOW), wherein standard lab chow was freely available in the operant chamber in weigh boats fastened to the chamber floor. This chow was weighed prior to and after each trial. The 30-minute trials were video recorded via USB cameras placed inside the sound-attenuating cubicles (Logitech HD 720P and Microsoft LifeCam Cinema). Time spent eating the freely available chow and total lever presses were recorded.

Outcome Devaluation: An outcome devaluation session was performed prior to PR/CHOW testing. Reinforcers and lab chow were devalued through refeeding, wherein all mice were given free home cage access to standard lab chow and reinforcer pellets 20 and 4 hours, respectively, prior to PR/CHOW testing. Time spent consuming chow and total lever presses were recorded.

Testing: Following outcome devaluation, mice underwent 4 days of PR/CHOW testing. Mice were again allowed to lever press at increasing increments for a reinforcer pellet or consume the standard lab chow freely available on the chamber floor. 30 minutes prior to PR/CHOW test trials, mice were injected i.p. with either CNO or Vehicle on alternating days. Treatment days were counterbalanced, so half of the mice received CNO on days 1 and 3 and Vehicle on days 2 and 4. Lever-pressing and time spent consuming chow were averaged, allowing comparisons between subjects according to virus and within subjects according to CNO treatment.

### Free Feed Test

CRF-ires-Cre mice were maintained at 85-90% of their ad libitum body weights for two days after the final day of concurrent choice testing and then completed two days of free feed testing, prior to which mice were injected i.p. with CNO and Vehicle on alternating days. Free feed testing was performed in the home cage with a pellet of standard lab chow for 1 hour. Chow consumed as a proportion of body weight was calculated.

### Open Field (OF) Test

Following the free feed test, CRF-ires-Cre mice were allowed ad libitum access to food and water for one week. OF testing was then performed in a standard Plexiglas open-field chamber (43 x 43 cm). Motor Monitor software (Kinder Scientific, Poway, CA, USA) recorded the mouse position throughout the test.

### Short-term Sucrose Preference Test

11 days after OF testing, CRF-ires-Cre mice were habituated to a two-bottle choice by placing both a 1% sucrose solution and standard drinking water in their home cage. After 48 hours of habituation, animals were fasted from both food and water for 18 hours before undergoing a short-term sucrose preference test, wherein all animals were injected with CNO 30 minutes prior to testing, then were returned to their home cages with two-bottle choice for a single one-hour sucrose preference test. One female mouse was removed from testing due to a decline in body weight below 85% of her ad libitum body weight. Bottles were weighed before and after testing, and volume of sucrose solution and regular drinking water consumed was recorded.

### Statistical Analysis

During effort-related choice, sucrose preference, and open field, all animals were injected with CNO. The effects of chemogenetic activation on effort-related choice was determined by repeated measures two-way ANOVAs with viral treatment with DREADD (male, *n* = 4; female, *n* = 6) or GFP control virus (male, *n* = 4; female, *n* = 4) as a between-subjects factor and barrier height as within-subjects factors. T-tests were used to compare the effects of DREADD virus (DREADD, *n* = 12) with control virus (mCherry, *n* = 9) on OF locomotion and time spent in the center of the OF. The effect of CNO administration on DREADD-free controls was assessed by two-way ANOVAs with sex and treatment with CNO (male, *n* = 6; female, *n* = 7) or Vehicle (male, *n* = 6; female, *n* = 7) as between-subjects variables, and barrier height as a within-subjects factor.

To assess sex differences in baseline PR and PR/CHOW training data, two-way ANOVAs with sex (male, *n* = 12; female, *n* = 9) as a between-subjects variable and training day or condition (PR, PR/CHOW, Re-fed) as within-subjects factors. Concurrent choice and free feeding tests used a within-subjects design by alternating CNO and vehicle administration in the same mice. The effect of chemogenetic activation was determined by two-way ANOVAs, with sex and viral treatment with DREADDs (male, *n* = 7; female, *n* = 5) or control mCherry (male, *n* = 5; female, *n* = 4) as between-subjects factors and CNO/Vehicle administration as a within-subjects factor. The effect of CNO administration on sucrose preference was assessed using a two-way ANOVA, with viral treatment as a between-subjects factor and sucrose/H2O consumption as a within-subjects factor. When main effects or interactions were found, Sidak’s post-hoc tests for multiple comparisons were performed.

## Results

### Chemogenetic activation of CRF+ aBNST neurons increases avoidance behavior in both male and female mice

We bilaterally injected aBNST of CRF-ires-Cre mice with AAV encoding Cre-dependent Gq (activating)-DREADD (n = 22) or Cre-dependent control (GFP/mCherry) virus (n = 17) (**Fig. 1A**). Viral expression endured throughout testing, and CRF+ neurons were observed throughout aBNST (**Fig. 1B**). Optogenetic or chemogenetic activation of these CRF+ neurons increase avoidance without affecting overall locomotion [15, 16, 26]. Similarly, we found that Gq-DREADD mediated activation (**Fig. 1C**) of aBNST CRF+ neurons increased avoidance of the center of an open field. Gq-DREADD-injected mice spent significantly less time in the center of the OF compared to mCherry controls (unpaired t-test, *t* (19) = 2.331, *p* < 0.05) (**Fig. 1D**). There was no effect of sex (*F*(1,17) = 4.092, *p* = 0.059) on time spent in the center (**Supp. 1A**), and no effect of chemogenetic activation on locomotion as distanced traveled in the OF was not statistically different between DREADD and mCherry control mice (*t* (19) = 0.242, *p* = 0.811) (**Supp. 1B**).

**Figure 1.**
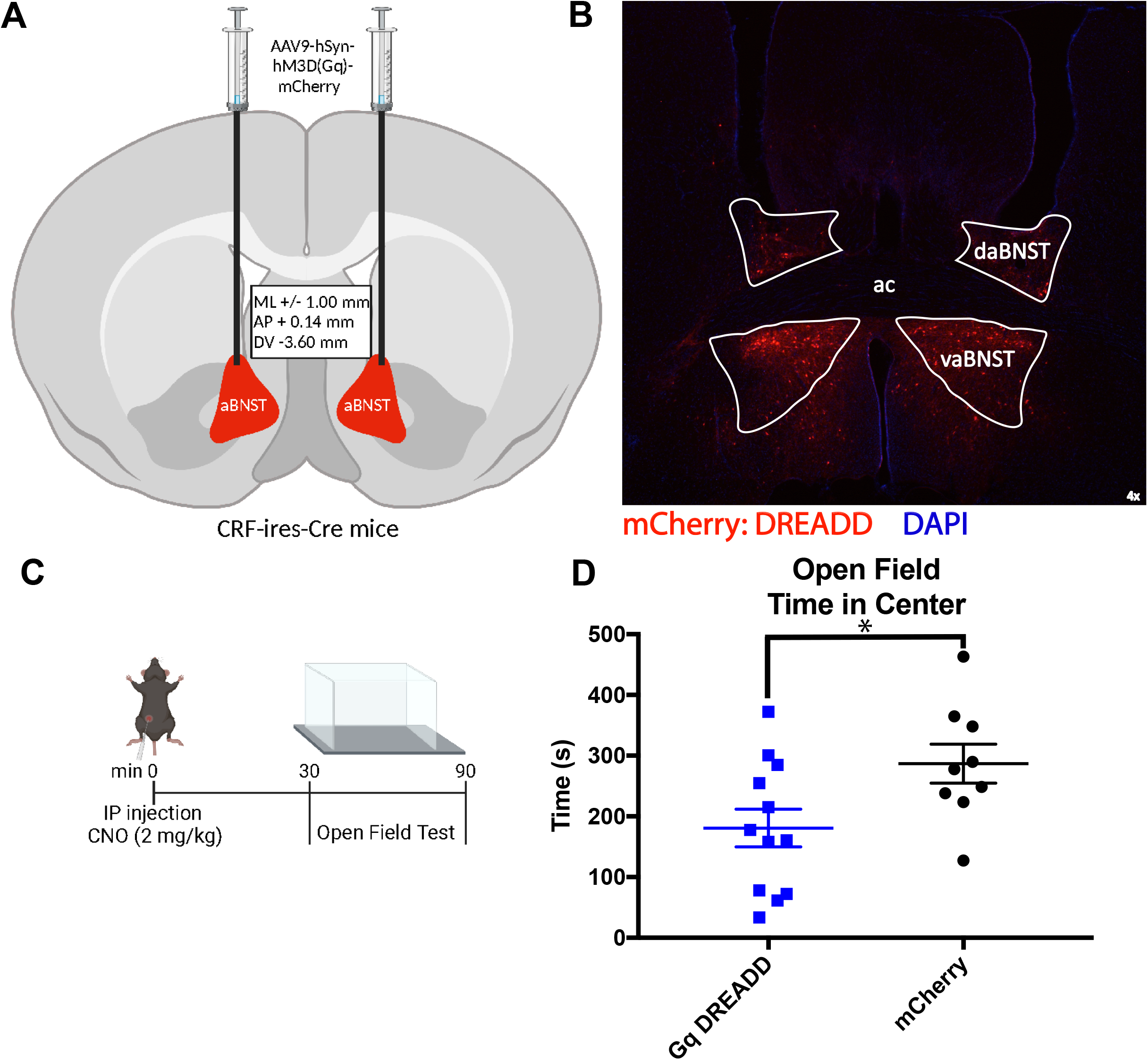
Virally-mediated DREADD expression was confirmed to localize in the anterior bed nucleus of the stria terminalis (aBNST), a region associated with avoidance behaviors. Viral expression was induced through (**A**) bilateral infusion of Cre-dependent DREADD virus at the anterior dorsolateral BNST of transgenic CRF-ires-Cre mice. Created with BioRender.com. (**B**) Viral expression, marked by red fluorescent marker mCherry, occurred throughout the aBNST. White lines delineate BNST borders of the dorsal aBNST, ventral aBNST (vaBNST) and the anterior commissure. Brain anatomy based on Allen Brain Atlas. (**C**) CNO was injected 30 minutes prior to testing. (**D**) As expected, chemogenetic activation of the CRF-expressing neurons in the anterior BNST via clozapine-N-oxide (CNO) injection increases avoidance behavior in the open field, reducing time spent in the center for Gq-DREADD (M = 180.8 s, blue squares) but not mCherry control (M = 286.7 s, black points) mice (*t* (19) = 2.331, *p* < 0.05). Values are plotted as individual values and mean +/- SEM. \**p* < 0.05

### Chemogenetic activation of CRF+ aBNST neurons in both female and male mice decreases willingness to exert effort for a high value reward in effort-related choice

Chronic stress leads to sustained increases in activity of CRF+ aBNST neurons [16, 19, 21–23] and impairs effortful motivation behavior [3, 5]. Therefore, we wondered whether chemogenetic activation of aBNST CRF neurons would be sufficient to mimic the effects of chronic stress on effort-related choice. To test the hypothesis, a cohort of CRF-ires-Cre mice received bilateral injections of AAV encoding Cre-dependent Gq-DREADD (*n* = 10; 6 female, 6 male) or GFP control (*n* = 8; 4 female, 4 male). Mice were allowed to recover from surgery for three weeks and then fasted to 85-90% of their ad libitum body weight. They then underwent training and testing in a Y-maze barrier task [3] to assess the effects of aBNST CRF+ neuron activation on effort-related choice (**Fig. 2A**). First, mice were trained in a Y-shaped maze to select a high reward (HR) arm containing 4 reinforcer pellets over a low reward (LR) arm containing 2 reinforcer pellets at the ends of the Y-maze arms. After 16 days of training, all mice reached a criterion of 70% HR arm choice: in at least 7 of the 10 daily free-choice trials subjects chose the HR arm (**Supp. 2A**).

**Figure 2.**
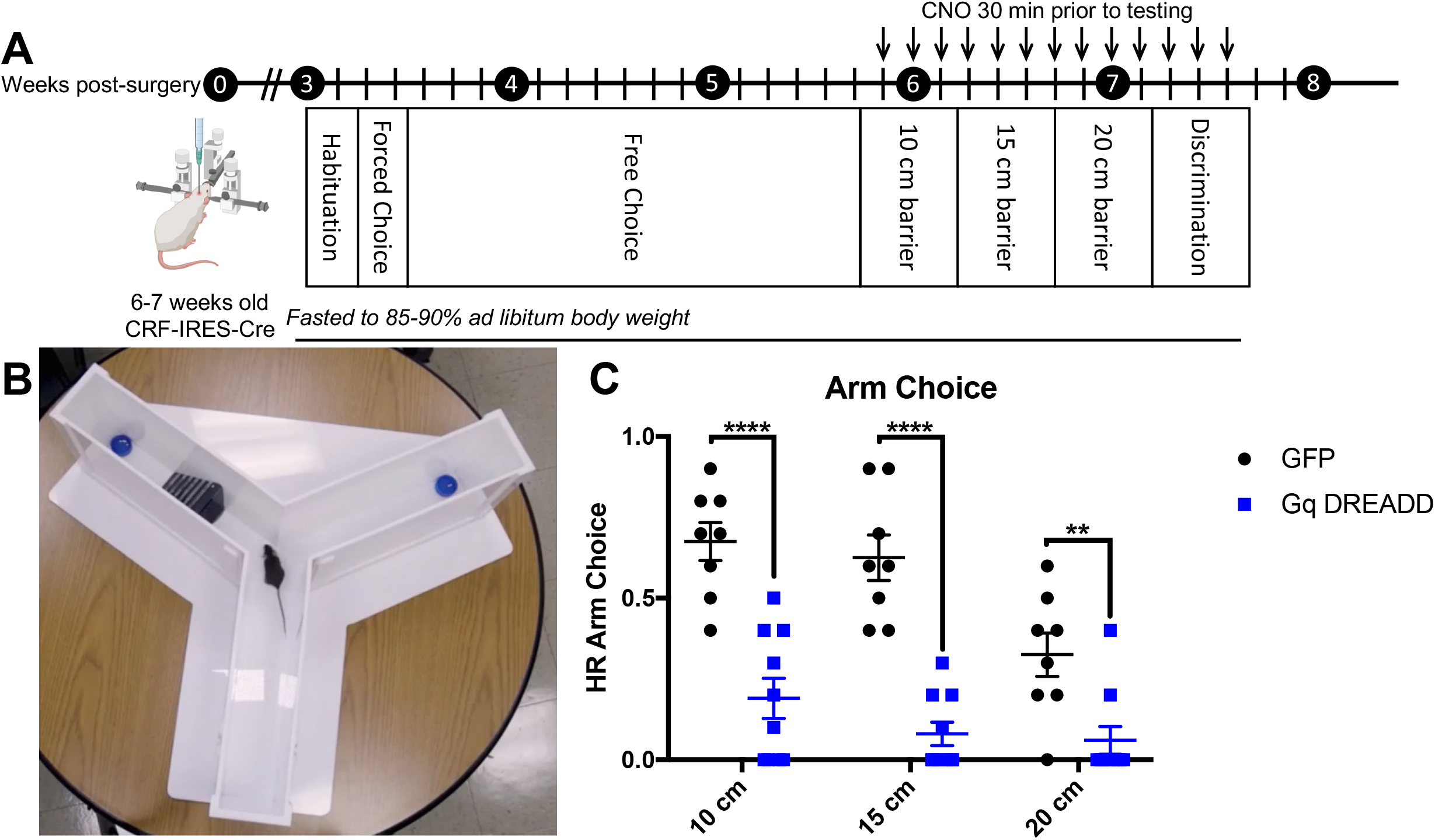
Chemogenetic activation of CRF+ BNST neurons reduced selection of the high reward (HR) arm in a Y-maze barrier task. (**A**) 6-7 week old CRF-ires-Cre mice were bilaterally injected with Cre-dependent Gq-DREADDs or a control Cre-dependent GFP virus. After recovery, mice were fasted and began training in the effort-related choice Y-maze barrier task. Training included 2 days of habituation, 2 days of forced chocie training, and 18 days of free choice training, wherein mice were trained in a Y-shaped maze to select a HR of of 4 sucrose pellets over a low reward (LR) of 2 pellets. Mice then underwent 3 days of 10 cm barrier testing, 3 days of 15 cm barrier testing, and 3 days of 20 cm barrier testing, prior to which mice were injected with CNO. During barrier testing, barriers were placed in the HR arm (**B**), so mice could choose to climb the barrier for an HR or traverse the other arm for an LR. (**C**) Gq-DREADD (blue squares) activation reduced HR arm choice, compared to GFP controls (black points), when tested in 10 (*p* < 0.001), 15 (*p* < 0.001) and 20 cm (*p* < 0.005) barrier conditions. HR arm choice is a proportion of 10 trials during which mice chose the HR arm. Values are plotted as individual values and mean +/- SEM. \*\**p* < 0.01; \*\*\*\**p* < 0.0001.

Next, a barrier (10, 15, or 20 cm) was placed in the HR arm (**Fig. 2B**). Having to climb this barrier increases the effort requirements for the higher value reward while the lower value reward remains freely available [41, 42]. Both female and male mice exposed to chronic stress shift to choosing the LR arm at shorter barrier heights than control mice [3, 5]. All mice were administered i.p. injection of 2 mg/kg CNO 30 minutes prior to testing, thereby activating CRF+ aBNST neurons in Gq-DREADD mice. The mice underwent 10 trials per day over 3 days each of 10, 15, and 20 cm barrier testing. A repeated measures two-way ANOVA with virus (DREADD or GFP) as a between-subjects factor and barrier height (10, 15, 20 cm) as a within-subjects factor demonstrated a significant interaction between virus and barrier height (*F*(2,32) = 6.24, *p* < 0.0051). The control mice chose the HR arm significantly more often than Gq-DREADD mice at all barrier heights, including 10 cm (*p* < 0.001), 15 cm (*p* < 0.001), and 20 cm (*p* < 0.005). As expected, all mice chose the HR arm significantly less when presented with a 20 cm tall barrier, compared to when presented with 10 (*p* < 0.0001) or 15 (*p* < 0.01) cm tall barriers (**Fig. 2C**). These results are from cohorts containing both female and male mice, as we saw no effect of sex at any barrier height (**Supp. 2B**). Therefore, highly specific Gq-DREADD-mediated activation of CRF+ aBNST neurons mimics the effects of stress on effort-related choice behavior.

Following 20cm barrier testing, we then placed 10 cm barriers in both arms for 3 days of discrimination testing. Surprisingly, a two-way ANOVA with virus treatment (DREADD or Control GFP) as a between-subjects factor and testing day (days 1, 2, and 3) as a within-subjects factor indicated a significant interaction between testing day and virus treatment (*F*(2,32) = 4.02, *p* <0.05) (**Supp. 2C**). Multiple comparisons indicate that DREADD mice chose the HR arm significantly less than GFP Control mice on days 2 (*p* < 0.001) and 3 (*p* < 0.0001). One possible explanation for this result is that Gq-DREADD mice chose the HR arm so infrequently over the previous 9 days of 10, 15, and 20 cm barrier testing that they became overtrained to the LR arm.

CNO can be converted to clozapine, which may lead to the behavioral effects seen in effort-related choice [43] [44]. To confirm that CNO itself does not alter behavior in the Y-maze barrier task, female (*n* = 14) and male (*n* = 12) WT C57BL6/J control mice underwent effort-related choice training and reached 70% HR arm choice after 16 days of training (**Supp. 3**). WT mice were then injected i.p. with either CNO (*n* = 13) or Vehicle (*n* = 13) 30 minutes prior to Y-maze barrier testing. A two-way ANOVA with treatment (CNO, Vehicle) as a between-subjects factor and barrier height as a within-subjects factor indicated a main effect of barrier height (*F*(2,48) = 12.58, *p* < 0.0001) but not treatment (*F*(1,24) = 1.05, *p* = 0.324). For all mice, HR arm choice was significantly lower when a 20 cm barrier was placed in the HR arm. Therefore, CNO has no effect on effort-related choice.

### Male mice lever press more than female mice during progressive ratio and concurrent choice testing

To evaluate the effect of chemogenetic activation of CRF+ aBNST neurons on motivated behavior in a concurrent choice test, CRF-ires-Cre mice underwent stereotaxic surgery and received bilateral injection of either Gq-DREADD virus (*n* = 12; 5 female, 7 male) or mCherry control virus (*n* = 9; 7 female, 5 male) into the aBNST. After recovery, mice underwent training and testing in an instrumental progressive ratio with concurrent chow (PR/CHOW) task (**Fig. 3A**). Mice first underwent training to promote lever pressing in operant chambers, beginning with two days of magazine training, then two days of a fixed ratio (FR) 1 reinforcement schedule training, followed by single-sessions of variable ratio (VR) 2, VR5, and VR20 training. Then, baseline progressive ratio (PR) measures were obtained via 10 daily 30-minute sessions, during which mice lever pressed on an FR(n) x 15 schedule, requiring mice to press at an incrementally increasing frequency every 15 reinforcer pellets (FR1×15, FR2×15, etc.) (**Fig. 3B**). A two-way ANOVA with sex as a between-subjects variable and training day (1-10) as a within-subjects variable indicates a main effect of sex (*F*(1,19) = 11.02, *p* < 0.05) on baseline PR lever pressing. Females lever pressed less than males throughout this progressive ratio task.

**Figure 3.**
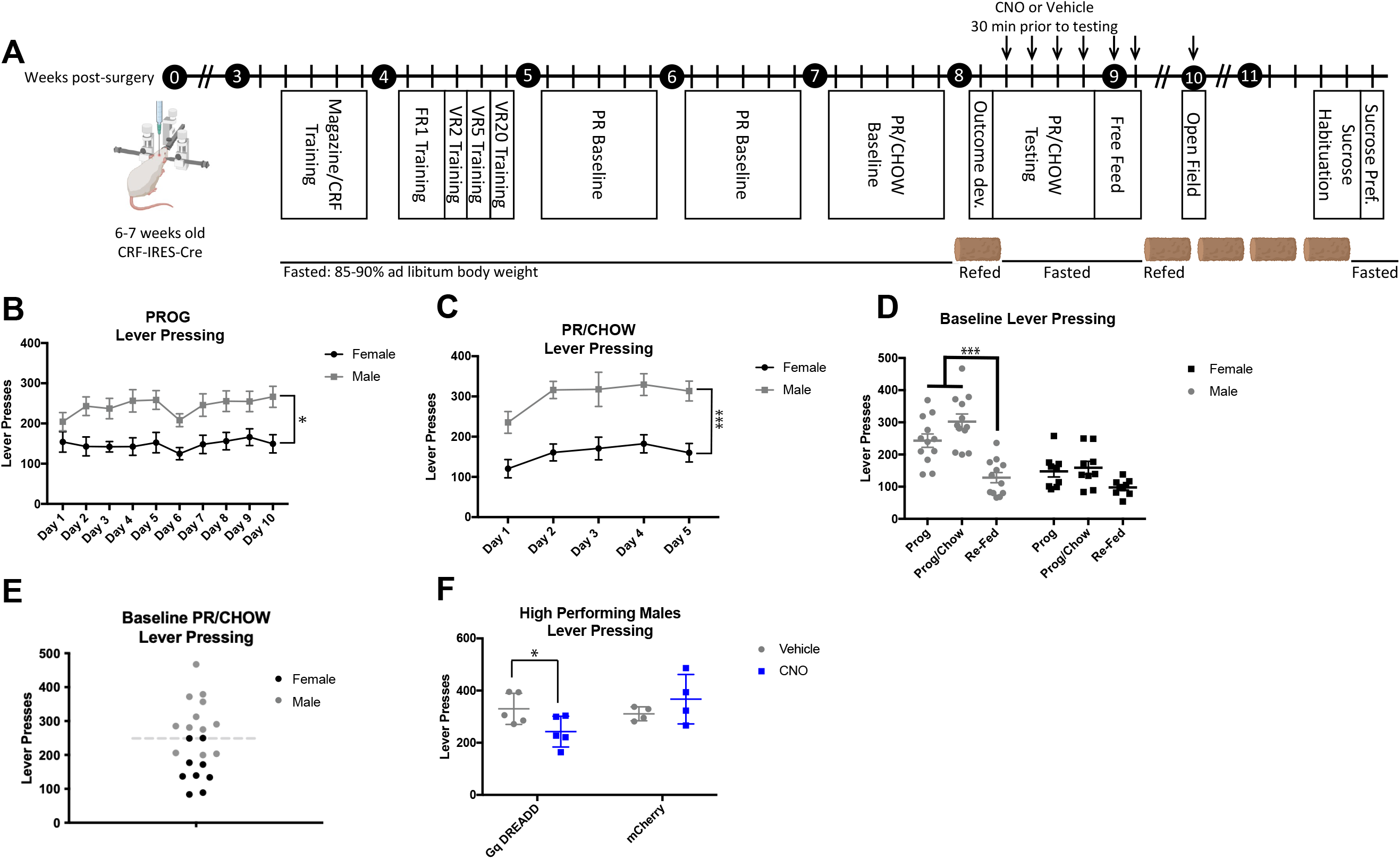
Chemogenetic activation of CRF+ BNST neurons decreased lever pressing for sucrose pellets in high-performing males. (**A**) 6–7-week-old CRF-ires-Cre transgenic mice were bilaterally injected with Cre-dependent Gq-DREADD or a control Cre-dependent mCherry virus. After recovery, mice were fasted and trained in operant chambers to lever press for sucrose pellets. Over 10 days, a progressive ratio (PR) baseline frequency of lever pressing was collected, wherein mice were trained to lever press at increasing frequencies in an (FR(n+1) x 15) PR reinforcement schedule. During PR/concurrent chow (CHOW) baseline training, concurrently available lab chow was freely available on the chamber floor. Mice were then refed and allowed ad libitum access to lab chow for 20 hours, and sucrose pellets for 4 hours, prior to an outcome devaluation trial. During 4 days of PR/CHOW testing, mice were injected with alternating treatments of CNO or Vehicle 30 minutes prior to PR/CHOW testing, to assess how chemogenetic activation of CRF+ BNST neurons alters lever pressing in the PR/CHOW task. Appetite was assessed over two days of free feed testing, wherein mice were injected with alternating treatments of CNO or Vehicle. During open field and sucrose preference testing, all mice were injected with CNO solution. Mice were fasted to 85-90% of their ad libitum body weights throughout training and testing, except where marked with a chow icon, which represents free access to standard lab chow. Arrows represent daily CNO or Vehicle injections, 30 minutes prior to testing. (**B**) Males (grey) lever pressed significantly more frequently than females (black) during PR baseline training (*p* < 0.05) on days 2, 4, 5, 7, 8 (*p* < 0.05), and 10 (*p* < 0.01). (**C**) Males also pressed more than females during PR/CHOW (*p* < 0.001) baseline training, including days 1 (*p* < 0.05), 2 (*p* < 0.001), and 3-5 (*p* < 0.01). (**D**) Males lever pressed significantly less when re-fed during outcome devaluation, compared to PR (*p* < 0.001) or PR/CHOW (*p* < 0.0001) baseline. Female lever pressing did not significantly decline during outcome devaluation. (**E**) Mice were designated high performers if their average PR/CHOW baseline lever pressing was greater than the median lever pressing (perforated line) for all mice. All high-performing mice were males (gray points), but all females (black points) were at or below the median baseline frequency. (**F**) In the PR/Chow test, high-performing Gq-DREADD males, when injected with CNO, lever pressed less than when injected with Vehicle (*p* < 0.05). No effect of CNO was observed in mCherry control mice (*p* = 0.884). Values are plotted as individual values and mean +/- SEM. \**p* < 0.05; \*\*\**p* < 0.001.

Baseline PR/CHOW lever pressing measures were then obtained over 5 days, during which mice could either exert effort by lever pressing on a PR schedule to consume a palatable reinforcer pellet or consume standard lab chow freely available on the chamber floor (**Fig. 3C**). A two-way ANOVA revealed a main effect of sex (*F*(1,19) = 19.93, *p* < 0.001). Females lever pressed less than males throughout this concurrent choice task.

Prior to PR/CHOW testing, one trial of outcome devaluation assessed lever pressing when mice were in a refed state. All mice were given access in their home cages to standard lab chow 20 hours prior to testing and reinforcers 4 hours prior to testing. A two-way ANOVA with sex as a between-subjects variable and condition (PR, PR/CHOW, Refed) as a within-subjects variable revealed a significant interaction between sex and condition (*F*(2,57) = 4.307, *p* < 0.05), with main effects of both testing condition (*F*(2,57) = 19.44, *p* < 0.0001) and sex (*F*(1,57) =, *p* < 0.0001). Multiple comparisons indicate that male, but not female, mice lever pressed significantly less in the refed state, compared to during fasted PR (*p* < 0.001) and fast PR/CHOW (*p* < 0.0001) (**Fig. 3D**). Therefore, the satiety-based outcome devaluation was effective in male mice.

Results from baseline PR, baseline PR/CHOW, and outcome devaluation suggested that individual differences in lever pressing may affect ability to detect reductions in motivation. Therefore, we separated mice into high and low performers. Previous studies [45, 46] also split subjects into high and low performers based on vehicle or baseline lever pressing. Average lever pressing over the 5 days of PR/CHOW baseline training was calculated, and a median lever pressing value was obtained. Subjects with average lever presses above the median were designated high performers, and those below the median were designated low performers (**Fig. 3E**). The divide between high-performing and low-performing mice was primarily driven by sex as no females were in the high-performing group. Since low-performing mice were not lever pressing enough to detect a possible Gq-DREADD mediated decrease in lever pressing, we focused our analyses on the high-performer male mice.

### Chemogenetic activation of CRF+ aBNST neurons decreases willingness to exert effort for a high value reward in a concurrent choice task in high-performing male mice

All high-performing male mice received CNO and Vehicle on alternating days of the 4 total days of PR/CHOW concurrent choice testing; thus, every mouse received two days of CNO injections and two days of Vehicle injections. Mice were injected i.p. 30 minutes prior to PR/CHOW testing and allowed to either lever press at a PR schedule or consume standard lab chow freely available on the chamber floor. A repeated measures two-way ANOVA of the high-performing male mice revealed a significant interaction between CNO treatment (CNO, VEH) and virus (Gq-DREADD, Control mCherry) (*F*(1,9) = 0.022, *p* < 0.05). Multiple comparisons established that Gq-DREADD mice lever pressed significantly less when injected with CNO compared to when they were injected with Vehicle (*p* < 0.05). CNO did not alter lever pressing in control mCherry mice (*p* = 0.884) (**Fig. 3F**). Therefore, Gq-DREADD mediated activation of CRF+ aBNST neurons in high-performing male mice decreased lever pressing for a high value reward when a lower value standard chow was freely available.

### Chemogenetic activation of CRF+ aBNST neurons does not decrease free feeding

To assess whether reductions in effortful behavior in effort-related choice or concurrent choice testing are due to decreased feeding behavior, CRF-ires-Cre mice underwent a two-day free feeding test 24 hours following PR/CHOW testing. Mice remained fasted to 90% of their ad libitum body weight and, 30 minutes prior to testing, were injected with CNO or Vehicle on alternating days. Mice were then placed in their home cage with a pellet of standard lab chow for 1 hour, and chow consumed as a proportion of body weight (Chow Consumed (g)/Body Weight (g)) was calculated. A two-way ANOVA revealed a significant interaction between virus treatment and CNO/VEH treatment (*F*(1,19) = 8.611, *p* < 0.01), but no main effect of CNO treatment alone (*F*(1,19) = 0.552, *p* = 0.467) or virus treatment alone (*F*(1,19) = 0.801, *p* = 0.382). Multiple comparisons found that DREADD mice injected with CNO actually consumed significantly more chow relative to their body weight compared to those injected with Vehicle in this free feeding task (*p* < 0.05) (**Fig. 4A**). A two-way ANOVA with treatment and sex as a between-subjects variables indicated a significant effect of sex on chow consumed relative to body weight (F(1,34) = 8.855, *p* < 0.01). Females at less than males, regardless of virus or CNO treatment (**Supp. 4A**). When free feeding analysis was performed in just high-performing males, we did not observe any effect of virus or CNO treatment (**Supp. 4B**). Overall, these results indicate that Gq-DREADD mediated activation of aBNST CRF+ neurons does not decrease free feeding.

**Figure 4.**
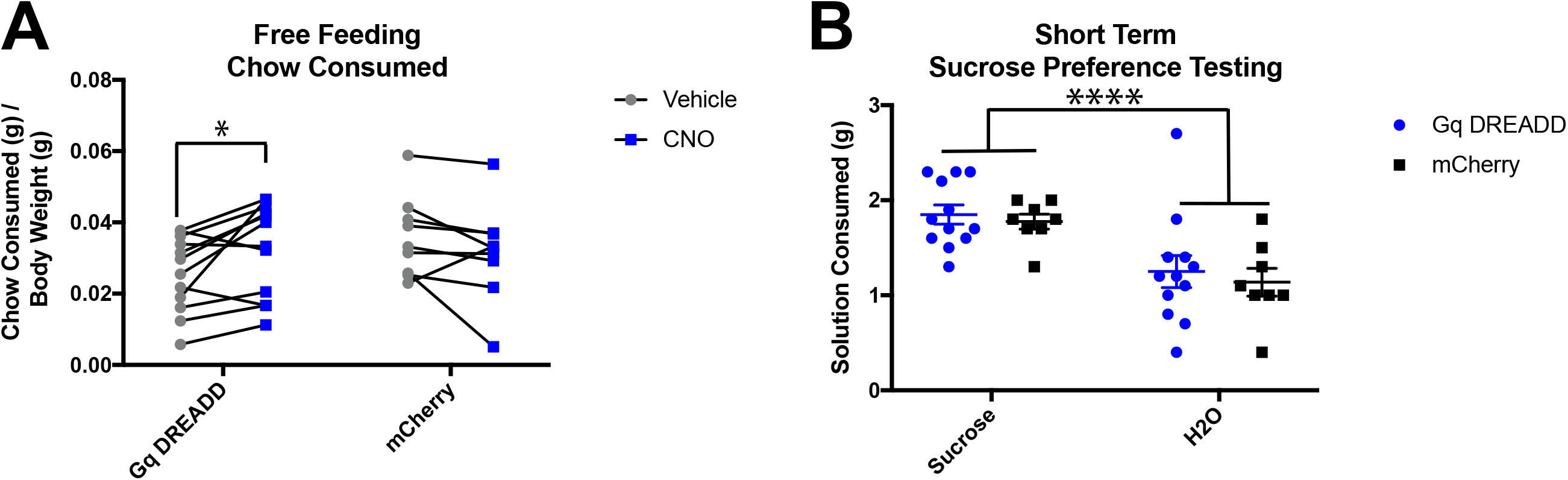
Chemogenetic activation of CRF+ BNST neurons increases appetite and does not affect sucrose preference. (**A**) Gq-DREADD and mCherry mice were injected with CNO or Vehicle over two alternating days, then chow consumption was measured in the home cage for 1 hour. Chow consumption in proportion to body weight was significantly higher in Gq-DREADD mice injected with CNO (blue points), compared to those same mice injected with Vehicle (gray points) (*p* < 0.05). (**B**) Sucrose preference was measured in mice deprived of food and water for 18 hours and injected with CNO 30 minutes before given the choice between 1% sucrose solution or water. All mice preferred sucrose solution over water (*p* < 0.01), and there was no effect of virus (*p* = 0.567) on sucrose consumption. Values are plotted as individual values and mean +/- SEM. \**p* < 0.05; \*\*\*\**p* < 0.0001. Taken together, appetite and sucrose preference do not explain the reduction in effort observed in the Y-maze barrier task or PR/CHOW.

### Chemogenetic activation of CRF+ aBNST neurons does not alter sucrose preference

To exclude reward preference as a factor in reducing lever pressing and barrier-climbing, a short-term sucrose preference protocol was performed after the free feeding and open field testing. For 48 hours, mice in their home cages were habituated to a two-bottle choice between water and a 1% sucrose solution. Following habituation, mice were fasted for 18 hours with no access to food or water. During short-term sucrose preference testing, all mice were administered CNO 30 minutes prior to testing, then mice were placed in their home cage with the two-bottle choice for 1 hour. A two-way ANOVA test with virus treatment as a between-subjects variable and solution type (water, sucrose) as a within-subjects variable established a main effect of solution-type (*F*(1,18) = 30.08, *p* < 0.0001) on consumption, but no effect of virus treatment (*F*(1,18) = 0.341, *p* = 0.567). Multiple comparisons demonstrate that both DREADD (*p* < 0.01) and mCherry control (*p* < 0.01) mice preferred sucrose solution over water (**Fig. 4B**). There was no effect of sex on sucrose preference (**Supp. 4C**), and when high-performing males were analyzed, there was no effect of virus or CNO treatment on sucrose preference (**Supp. 4D**). Therefore, Gq-DREADD mediated activation of aBNST CRF+ neurons does not alter reward preference.

## Discussion

Exposure to chronic stress increases avoidance behaviors and decreases motivation behaviors. aBNST CRF+ neurons are highly implicated in stress-induced avoidance [16, 19, 21, 22, 39], but it was previously unknown that they also play a role in effortful responding. A potential mechanism is that stress drives drug-seeking [31] and activates CRF+ BNST projections that inhibit dopaminergic reward-related regions, including the VTA and NAc [30, 33], thereby reducing motivated behavior. Here, we tested whether highly specific chemogenetic activation of CRF+ aBNST neurons mimics the effects of chronic stress on two tasks that assess willingness to exert effort for a high value reward when a low value reward is freely available: effort-related choice and concurrent choice.

The Y-maze barrier task used for effort-related choice [3] was adapted from Salamone et al. [47] in order to assess cost/benefit analysis in mice. Chemogenetic activation of CRF+ aBNST neurons resulted in both male and female mice choosing the low reward arm more often than control mice at all barrier heights. These results demonstrate that activating CRF+ aBNST neurons reduces effort-related choice. This reduction in HR arm choice is similar to what is observed in male and female mice exposed to chronic stress and in male mice exposed to chronic corticosterone administration [3–5]. Taken together, these results suggest that specific activation of CRF+ aBNST neurons is sufficient to mimic the effects of chronic stress on effort-related choice.

After effort-related choice testing was completed, we exposed mice to three days of discrimination testing, where 10 cm barriers were placed in both the HR and LR arms. Surprisingly, Gq-DREADD mice continued to choose the LR arm during discrimination. This arguably could be a learning deficit, however, lesion studies suggest that the BNST is not necessary for spatial learning [48]. An alternative hypothesis is that the Gq-DREADD mice became overtrained after repeatedly choosing the LR arm over the 9 days of barrier testing, amounting 90 total trials. They failed to reverse their arm choice as a function of habitual behavior, wherein choosing the LR arm became automatic and resistant to change. Tendencies to repeat behaviors can be augmented by stress [4]. WT DREADD-free controls validated findings from the Y-maze barrier task, confirming that CNO alone does not alter arm choice and barrier climbing abilities in males nor females.

We also tested whether chemogenetic activation of CRF+ aBNST neurons decreased willingness to exert effort for high value reward using concurrent choice. Mice were presented with a progressively higher-effort/high-reward choice to lever press on a progressive ratio schedule for a palatable reinforcer pellet or a low-effort/low-reward choice to consume standard lab chow freely available on the chamber floor. The concurrent choice paradigm has been used to measure motivated behavior [49], and is largely associated with dopamine signaling and activity in mesolimbic reward-related brain regions, as reviewed by Salamone et al. in 2016 [50]. Baseline progressive ratio established a clear baseline sex difference, where female mice lever pressed significantly less than males in both PR and PR/CHOW conditions. Therefore, a floor effect existed in the females where it was difficult to see decreased lever pressing from a low baseline in both an outcome devaluation task and with Gq-DREADD mediated activation of CRF+ aBNST neurons in concurrent choice. Sex differences have been observed previously in appetitive learning and lever pressing tasks, where females are less engaged compared to males [51–54]. Future experiments are needed to determine the cause of these sex differences in lever pressing.

Previous studies [45, 46] have segregated rodents into high and low performers in PR/CHOW [45, 46, 56]. Using this previously published strategy, we divided mice using a median-split into high- and low-performers. All of the high-performers were males and most of the low-performers were females. During concurrent choice testing, high-performing male mice received injections of either CNO or Vehicle on alternating days, allowing treatment-dependent within-subjects comparisons. Experimental male mice lever pressed significantly less during DREADD-mediated activation of CRF+ aBNST neurons, compared to after receiving vehicle solution. This reduction in lever pressing is similar to what is observed in mice after exposure to chronic stress or corticosterone administration [3–5]. Taken together, these results suggests that activation of CRF+ aBNST neurons is sufficient to mimic the effects of chronic stress on concurrent choice testing in high-performing male mice.

### Role of CRF+ aBNST Neurons in Motivated Behavior

Experiments over the past decade that support a “valence surveillance” [57] role of the BNST used self-stimulation and real-time place preference (RTTP) tests to show that stimulation of specific BNST circuits [38] and cell types [15, 38] is rewarding. Activation of dorsal BNST neurons, including those that project to the paraventricular nucleus, promoted reward responding in a Pavlovian reward task [58]. However, RTPP, self-stimulation, and reward learning tests do not assess effortful behavior. Baumgartner et al. [37] found that rats lever pressed less when sucrose rewards were paired with optogenetic activation of CRF+ BNST, concluding that CRF+ BNST neurons promote aversive motivation-the drive to avoid negatively valenced states. This concurs with findings that CRF+ BNST neurons drive avoidance in aversive or threatening environments [15–17]. The role of the BNST in aversive motivation may also coincide with findings that CRF+ BNST neurons promote drug-seeking behaviors [35, 36], possibly to avoid withdrawal [59]. Our experiments are novel in that they assess a potential role of CRF+ BNST neurons in underlying the effects of stress exposure on incentive motivation – the drive to approach and work for baseline rewards, such as food.

Motivation behavior is highly associated with dopaminergic mesolimbic pathways. Projection-specific optogenetic stimulation of all VTA-projecting GABAergic BNST neurons is anxiolytic and rewarding [38]. Here, however, we used cell-specific chemogenetic manipulations that activated CRF+ aBNST neurons, which inhibit [17, 30] afferents via corelease of CRF and GABA [60, 61]. CRF+ BNST neurons project to the VTA [17, 27, 29, 30] and NAc [17], but the effects of CRF in these mesolimbic reward areas are complex [62]. Intra-VTA infusions of CRF reduce dopamine signaling and reduce motivated behavior in instrumental PR testing [33]. However, CRF infusion has also been shown to be rewarding [63], increasing dopamine signaling in the VTA [63] and driving motivated drug-seeking behavior [32]. These opposing results may be explained by a potential “valence flip”, wherein the effects of CRF depend on environmental conditions, such as substance abuse or stress [62, 64]. We hypothesize that the reduction in effortful motivation behaviors was driven by GABAergic inhibition of VTA and/or NAc by CRF+ aBNST neurons. Further experiments should investigate whether chemogenetic activation of CRF+ aBNST neurons results in co-release of CRF, and the nature of the modulatory role of CRF in mesolimbic reward areas. Experiments should also specifically target aBNST^CRF^-VTA and aBNST^CRF^-NAc projections in behaving animals.

### Sex Differences in CRF-Signaling

CRF signaling in the BNST differs between sexes [65–67]. We used both male and female mice to determine if activation of CRF+ aBNST neurons alters effortful motivation behavior differently across sexes. We did not see any effect of sex in effort-related choice. However, we cannot rule out that a statistical floor effect may have masked sex differences, because all mice expressing DREADDs almost completely ceased to choose the HR arm during testing. Sex differences were already evident during baseline concurrent chow training; thus, the effect of CRF+ aBNST neuron activation during testing could not be accurately compared between sexes. Future experiments will need to improve female engagement in the concurrent chow test, or design alternative tasks where both sexes can be tested.

### Conclusions

We used chemogenetics to mimic the activating effects of chronic stress on CRF+ aBNST neurons [16, 19, 21, 22, 39], and results collected from both sexes in effort-related choice and from high-performing males in the concurrent chow task demonstrate that chemogenetic activation of CRF+ aBNST neurons reduces effortful motivation behavior without affecting appetite, sucrose preference, or locomotion. These results demonstrate a critical role of CRF+ aBNST neurons in mediating effortful motivation behaviors.

**Table 1.** Experimental design of the Y-maze barrier task and PR/CHOW experiments.

| **Table 1** |  |  |  |  |  |  |
| --- | --- | --- | --- | --- | --- | --- |
|  | Mice | # Experimental | Experimental Group | # Control | Control Group | CNO |
| **Effort-Related Choice** | CRF-ires-Cre | 10 mice (4 M, 6 F) | Cre-dependent Gq-DREADD | 8 mice (4 M, 4 F) | Cre-dependent mCherry virus | All |
| **Effort-Related Choice** | Wildtype | 13 mice (6 M, 7 F) | CNO | 13 mice (6 M, 7 F) | Vehicle | 50% |
| **PR/CHOW** | CRF-ires-Cre | 12 mice (7 M, 5 F) | Cre-dependent Gq-DREADD | 9 mice (5 M, 7 F) | Cre-dependent GFP virus | All (on alternating days) |
| **Open Field** | CRF-ires-Cre | 12 mice (7 M, 5 F) | Cre-dependent Gq-DREADD | 9 mice (5 M, 7 F) | Cre-dependent GFP virus | All |
| **Free Feeding** | CRF-ires-Cre | 12 mice (7 M, 5 F) | Cre-dependent Gq-DREADD | 9 mice (5 M, 7 F) | Cre-dependent GFP virus | All (on alternating days) |
| **Sucrose Preference** | CRF-ires-Cre | 12 mice (7 M, 5 F) | Cre-dependent Gq-DREADD | 9 mice (5 M, 7 F) | Cre-dependent GFP virus | All |

