## Supplemental Figures for "Chemogenetic activation of corticotropin-releasing factor-expressing neurons in the anterior bed nucleus of the stria terminalis reduces effortful motivation behaviors"

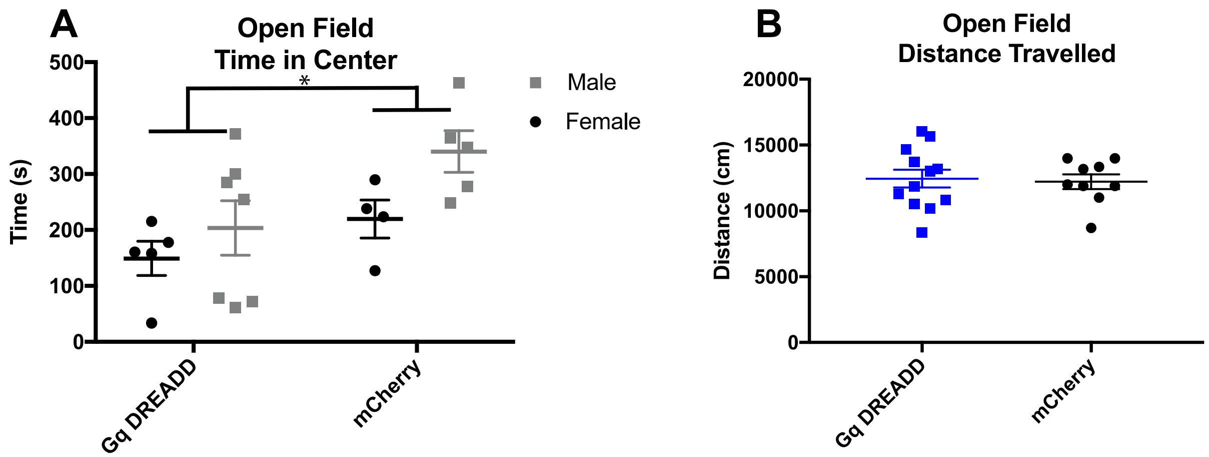


**Supplemental Figure 1.** Locomotor behavior (**A**) was not altered by chemogenetic activation of CRF-expressing BNST neurons, Gq-DREADD (M = 12,444 cm) and mCherry (M = 12,221 cm) mice traveled similar distances during the 30-minute OF test ($t$ (19) = 0.242, $p$ = 0.811). (**B**)No effects of sex on avoidance behavior in the open field (OF) test. Over the 30-minute OF test, time spent in the center of the OF did not differ between male (gray) and female (black) mice ($p$ = 0.059). Values are plotted as individual values and mean +/- SEM. *$p$ < 0.05.

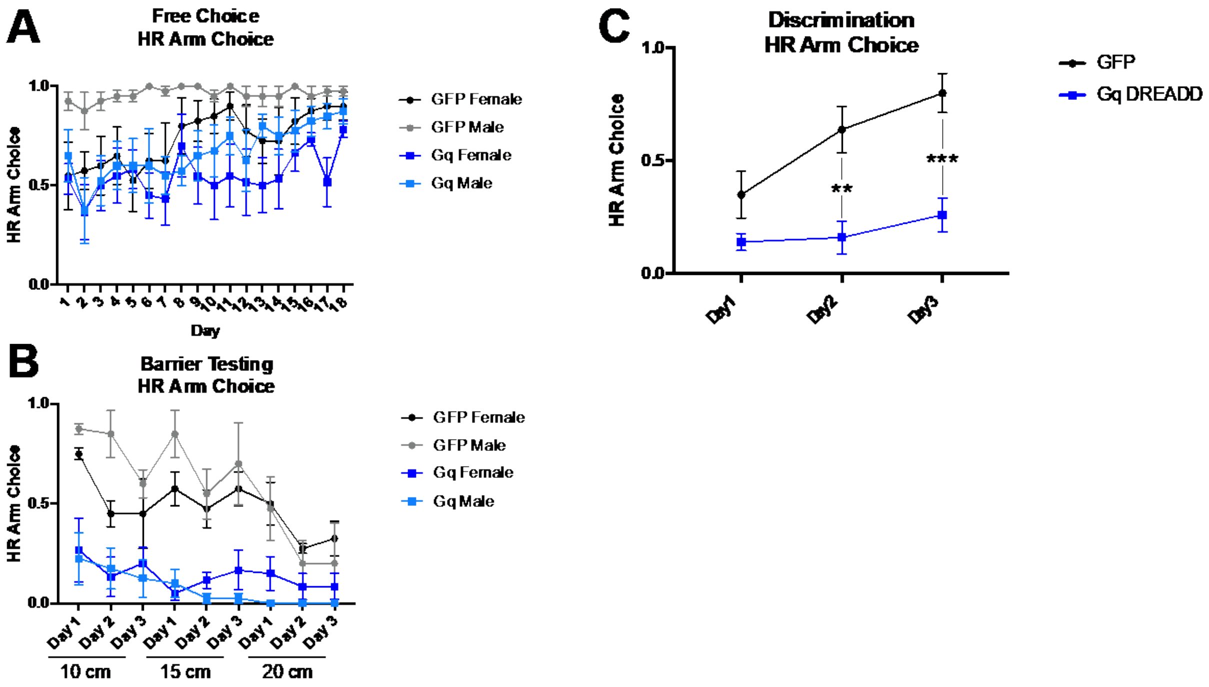


**Supplemental Figure 2.** CRF-ires-Cre transgenic mice learned to select the high reward (HR) arm during Y-maze training but failed to choose the HR arm during discrimination. (**A**) During free choice training, mice were allowed to choose between an HR of 4 pellets or a low reward (LR) of 2 pellets. HR arm choice is plotted as the proportion of the 10 daily trials in which mice chose the HR arm. By day 18, all mice reached criteria of at least 70% HR arm choice, wherein they chose the HR arm at least 7 of the 10 daily free choice trials. Mice were assigned to the CNO (dark or light blue squares) or Vehicle (black or gray points) group after free choice training. (**B**) During barrier testing, a barrier was placed mid-way down the HR arm and all mice were injected with clozapine-N-oxide (CNO). There was no effect of sex on HR arm choice over the 3 days of 10 cm barrier testing ($p$ = 0.331), 3 days of 15 cm barrier testing ($p$ = 0.514), or 3 days of 20 cm barrier testing ($p$ = 0.364). Values are plotted as mean HR arm choice over 10 daily trials. (**C**) Barrier testing was followed by discrimination testing, wherein a 10 cm barrier was placed in both the HR and LR arms. Over 3 days of discrimination testing, the average HR arm choice was greater for GFP control mice compared to Gq-DREADD mice on days 2 ($p$ < 0.001) and 3 ($p$ < 0.0001). Values are plotted as individual values and mean +/- SEM. **$p$ < 0.001; *** $p$ < 0.001

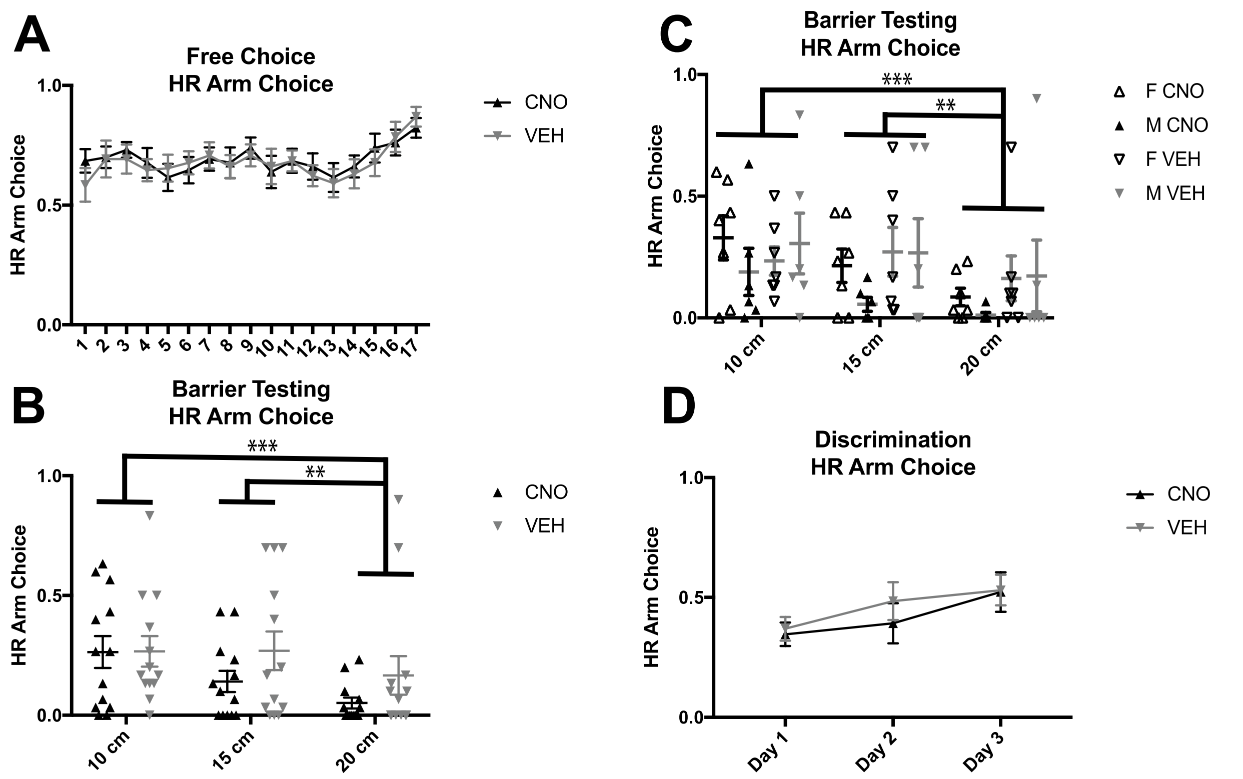


**Supplemental Figure 3**. No effect of CNO on the Y-maze barrier effort-related choice task in wildtype mice. (**A**) During free choice training, mice were allowed to choose between an HR of 4 pellets or a low reward (LR) of 2 pellets. By day 17, all mice reached criteria of at least 70% HR arm choice, wherein they chose the HR arm at least 7 of the 10 daily free choice trials. Mice were assigned to the CNO (black triangles) or Vehicle (gray triangles) treatment group after free choice training. Values are plotted as mean HR arm choice over 10 daily trials (**B**) During barrier testing, a barrier was placed mid-way down the HR arm and mice were injected with either CNO or Vehicle. There was no effect of CNO on HR arm choice ($p$ = 0.324). All mice chose the HR arm more often when the barrier was 10 cm ($p$ < 0.0001) and 15 cm ($p$ < 0.05) high, compared to when the barrier was 20 cm high. (**C**) Males and females chose the HR arm at similar rates during barrier testing, and there was no effect of sex on HR arm choice at 10 ($F$(1,22) = 0.131, $p$ = 0.721), 15 ($F$(1,22) = 0.770, $p$ = 0.390) or 20 cm ($F$(1,22) = 0.138, $p$ = 0.714). (**D**) Barrier testing was followed by discrimination testing, wherein a 10 cm barrier was placed in both the HR and LR arms. A two-way ANOVA indicated that mice injected with CNO or Vehicle chose the HR arm at similar rates ($F$(1,24) = 0.247, $p$ = 0.624). However, a main effect of testing day was revealed ($F$(2,48) = 6.735, $p$ < 0.01). Performance improved over the three days of discrimination testing, so mice chose the HR arm more often on day 3 compared to day 1 ($p$ < 0.01). Values are plotted as individual values and mean +/- SEM. **$p$ < 0.01; ***$p$ < 0.001.

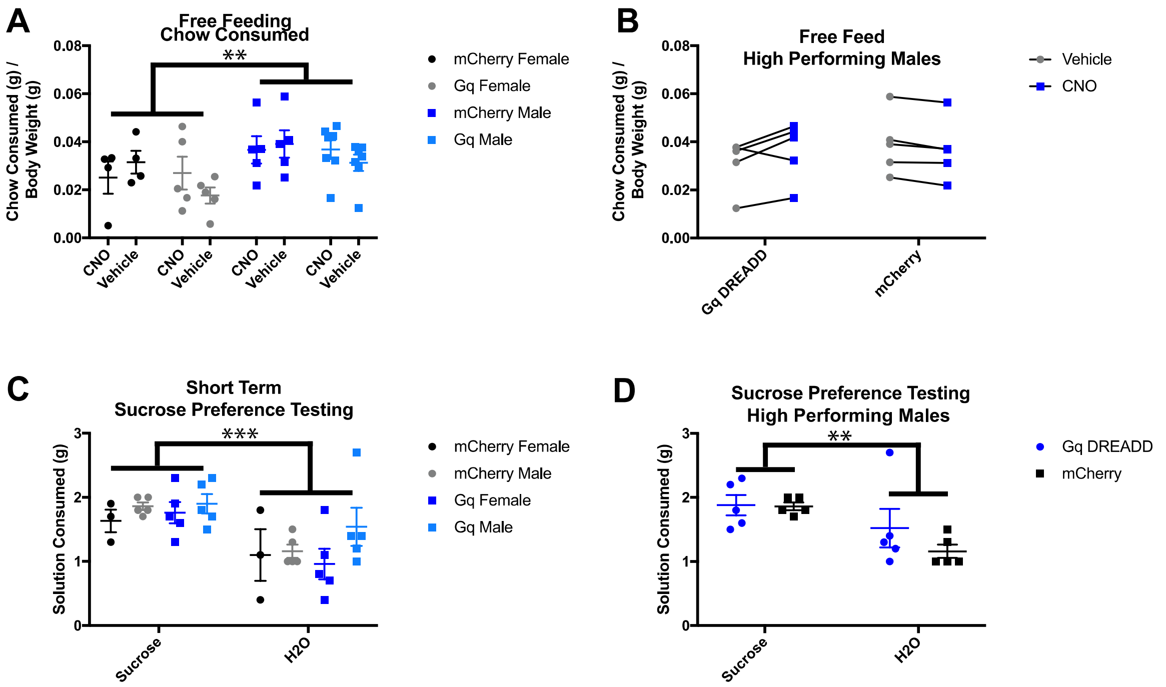


**Supplemental Figure 4.** Effects of sex and PR/Chow performance on behavior in free feed and sucrose preference tasks. (**A**) Gq-DREADD and mCherry male and female mice were injected with CNO or Vehicle over two alternating days, then chow consumption was measured in the home cage over 1 hour. Chow consumption in proportion to body weight was significantly higher in Gq-DREADD mice injected with CNO compared to those same mice injected with Vehicle ($p < 0.05$). Two-way ANOVA suggests sex differences in chow consumption ($p < 0.01$), but multiple comparisons do not indicate any one treatment group drives these sex differences. (**B**) Two-way ANOVA of only high-performing males indicates a significant interaction between virus treatment and CNO/Vehicle treatment, however, multiple comparisons between and within mice were not significant. (**C**) Sucrose preference was measured in mice deprived of food and water for 18 hours and injected with CNO 30 minutes before given the choice between 1% sucrose solution or water. Male and female mice preferred sucrose solution over water, and there was no effect of sex on sucrose preference. (**D**) Two-way ANOVA of only high-performing males indicates no effect of virus treatment on sucrose preference.
